# Deep Learning Restores Speech Intelligibility in Multi-Talker Interference for Cochlear Implant Users

**DOI:** 10.1101/2022.08.25.504678

**Authors:** Agudemu Borjigin, Kostas Kokkinakis, Hari M. Bharadwaj, Joshua S. Stohl

## Abstract

Cochlear implants (CIs) fail to provide the same level of benefit in noisy settings as in quiet. Current noise reduction solutions in hearing aids and CIs only remove predictable, stationary noise, and are ineffective against realistic, non-stationary noise such as multi-talker interference. Recent developments in deep neural network (DNN) models have achieved noteworthy performance in speech enhancement and separation. However, little work has investigated the potential of DNN models in removing multi-talker interference. The research in this regard is even more scarce for CIs. Here, we implemented two DNN models that are well suited for applications in speech audio processing, including (1) recurrent neural network (RNN) and (2) SepFormer. The models were trained with a dataset developed in house (*∼* 30 hours), and then tested with thirteen CI listeners. Both RNN and SepFormer models significantly improved CI listener’s speech intelligibility in noise without compromising the perceived quality of speech. These models not only increased the intelligibility in stationary non-speech noise, but also introduced substantially ***more*** improvements in non-stationary ***speech*** noise, where conventional signal processing strategies fall short with little benefits. These results show the promise of using DNN models as a solution for listening challenges in multi-talker noise interference.

## 1 Introduction

Cochlear implant (CI) users typically achieve satisfactory speech intelligibility in quiet environments such as audiology clinics, but struggle to converse in noisy social settings. There have been significant efforts towards the development of noise reduction algorithms in the field of hearing aids, and these algorithms are increasingly being translated into both CI research and commercial CI products. Noise reduction algorithms can be classified into single-vs multi-microphone strategies, as reviewed by Kokkinakis et al. (2012) and Henry et al. (2023).

A single-microphone noise reduction strategy involves processing the monophonic audio signal from a single microphone to reduce background noise. This can be achieved by using many approaches such as spectral subtraction (Yang & Fu, 2005; Verschuur et al., 2013), subspace estimation (Loizou et al., 2005) and gain adjustments based on the short-time signal-to-noise ratio (SNR) estimate (Mauger, Arora, & Dawson, 2012; Mauger, Dawson, & Hersbach, 2012; Cohen, 2003; Dawson et al., 2011) and ideal binary mask estimate (Hu & Loizou, 2008b; Koning et al., 2015). These traditional single-microphone algorithms are heavily driven by signal processing strategies based on time-averaged signal statistics and certain assumptions about the noise. For example, in spectral subtraction, the noise is assumed to be non-modulated and stationary (as well as broadband and additive), whereas the target speech signal can be retained by preserving the channels with relatively large amplitude modulations. Because of these inherent assumptions, these algorithms tend to perform relatively well only if the background is statistically consistent and predictable, such as a stationary noise. Yang & Fu (2005) observed improvement in speech recognition with speech-shaped noise but only moderate improvement with 6-talker babble. CI listeners in Mauger, Arora, & Dawson (2012)’s study showed improvements of 19% on average on speech recognition in speech-weighted noise and small improvement of 7% in 20-talker babble but no significant improvement in a 4-taller babble. Dawson et al. (2011) also observed greatest benefit in speech-weighted noise, compared to more realistic street-side city noise and cocktail party noise.

A number of studies evaluated a commercial noise reduction strategy—Advanced Bionic’s ClearVoice^TM^ (AdvancedBionics, 2012), which works by adjusting the short-term channel gains according to the estimate of instantaneous SNR. Dingemanse & Goedegebure (2015) did not see improvements in speech recognition with ClearVoice in steady state speech-spectrum noise. ClearVoice was only shown to increase performance in non-speech noise when combined with other parameter adjustments, such as adjusting maximum comfort levels (Dingemanse & Goedegebure, 2018), or with other technologies such as a remote microphone (Wolfe et al., 2015) and/or multi-microphone noise reduction strategies (Geißler et al., 2015; Hersbach et al., 2012). Similar commercial noise reduction strategies for CIs include the Ambient Noise Reduction (ANR) strategy from MED-EL (2021) and SNR-NR from Cochlear Limited (Mauger et al., 2014). Listening with CIs and other hearing assistive devices is especially challenging in scenarios with non-stationary noise, such as multi-talker background interference (Cullington & Zeng, 2008; Fu et al., 1998). Therefore, the inherent inability of single-channel noise reduction solutions to effectively remove non-stationary noise interference remains to be the most critical limitation of modern commercial noise reduction strategies for CIs.

Compared to single-microphone algorithms, multi-microphone noise reduction strategies, also commonly referred to as beamforming, involves capturing the audio signals from two relatively closely-spaced microphones and then processing the signals to reduce background noise from a spatial region, typically behind the listener. Beamforming was first introduced to commercial CI products by Cochlear Limited in 2005, over a decade after it became a standard feature in hearing aids (Kates & Weiss, 1996; Buechner et al., 2014). Beamforming typically consists of a front and rear microphone. A sound originating from behind arrives at the rear microphone before the front microphone. This external time delay is used by the signal processing to attenuate the sound from the back. Adaptive beamforming was later introduced to account for moving noise sources that are not always at the back of the listener but could be slightly off to the side from behind (Bentler et al., 2006). Directionality, the ability to focus on the signal in front of the listener, is typically better with more microphones (Dillon, 2008). However, due to the minimum distance requirement between the microphones and the size of a CI audio processor, it is impractical to have too many physical microphones on a single processor. Binaural beamformers were developed to increase the number of microphone signals available for beamforming by utilizing all four physical microphones from two hearing aids or audio processors for bilateral CI listeners. Picou et al. (2014) showed that binaural beamformer (4 microphones within 2 hearing aids) provided significantly better benefit in speech-in-noise hearing than monaural directional microphones (2 microphones within 1 hearing aid) regardless of the SNR level. Although multi-microphone noise reduction approaches have generally shown improvement in speech intelligibility in noise among CI listeners compared to conditions without any noise-reduction processing (Kokkinakis & Loizou, 2010), they require additional hardware and can be more complex to implement. The effectiveness of multi-microphone noise reduction strategies is limited to the scenario where the target speech and noise are spatially separated and the listener faces the target all the time. These algorithms can be adversely affected by reverberation, which exists in most listening situations, and can degrade source localization (Dawson et al., 2011; Henry et al., 2023). The effectiveness of a beamformer is also affected by how the device is worn by the listener. For a behind-the ear processor, the optimal sitting position of the device on the ear results in the front and rear microphones positioned close to horizontally, which allows for a match between the real delay and the assumed external time delay based on the physical distance between two microphones. This match enables relatively precise cancellation of noise. If the device is worn at an angle, for example, the real time delay of sound traveling from one microphone to the other will be smaller than the delay assumed by the strategy (Ricketts, 2001). Considering these restrictions from multi-microphone noise reduction solutions and the limitation of existing single-microphone algorithms in reducing non-stationary noise, we are interested in single-microphone noise reduction strategies that can suppress non-stationary, multi-talker noise interference.

Machine learning, especially deep learning, has transformed many fields, including audio processing. Machine learning appears to have the potential to overcome the limitation of current noise reduction solutions in CIs and hearing aids in reducing non-stationary noise (Henry et al., 2023). Research in machine learning-based noise reduction algorithms have only begun to surface in a larger scale in the last decade with the advent of more powerful computing resources and the availability of larger datasets. Early work took a small step forward by developing machine learning-based models for suppressing non-stationary, non-speech environmental noise and demonstrated improvement in objective quality measures (Gopalakrishna et al., 2012) and speech intelligibility among CI listeners (Hu & Loizou, 2010). More recent studies have shown improved speech intelligibility in non-stationary background noise such as a multi-talker babble. For example, Lai et al. (2017) showed improved speech intelligibility in 2-talker babble noise among normal hearing (NH) listeners with CI simulations with a deep denoising autoencoder (DDAE) compared to conventional noise-reduction approaches, and further improved the performance among Mandarin-speaking CI listeners when a noise classifier was added to the DDAE model (Lai et al., 2018). Goehring et al. (2017) demonstrated improved speech intelligibility in 20-talker babble among CI listeners when a neural network model was used to process the noisy speech.

Note that the models mentioned above operate based on *a priori* knowledge of the target and/or background by using the same materials from training during testing. For example, Hu & Loizou (2010), Goehring et al. (2017) and Lai et al. (2018) split the same speech corpus into training and testing datasets and used the same noise types to create noisy speech. The model performance is expected to decrease significantly if truly *unseen* testing data are presented to the models. A larger and more diverse dataset is key to improving model generalizability.

Another important aspect of generalizability is model architecture. Recent studies have demonstrated the potential of using recurrent neural network (RNN) models for better generalization by including a classic architecture called long short-term memory (LSTM) (Hochreiter & Schmidhuber, 1997). LSTM accumulates information from the past and hence enables the network to form a temporary memory, which is essential for properly managing and learning speech context. Many studies have demonstrated the success of RNN-LSTM based models in speech recognition, enhancement, and separation applications (Graves et al., 2013; Weninger et al., 2015; Chen & Wang, 2017; Kolbæk et al., 2017). However, the application of such models in noise reduction for hearing aids and CIs has been limited except for a few recent studies, such as Healy et al. (2019) and Goehring et al. (2019).

The present work aims to further explore the potential of machine learning models in reducing non-stationary, multi-talker noise interference. We also aim to develop models with greater generalizability by leveraging a large training dataset as well as advanced model architectures, such as RNN. Despite the wide adoption of RNN in modern audio processing systems and in many other domains, RNN architecture suffers from “vanishing gradient” or “short-term memory”, which renders them ineffective for long speech instances. Therefore, in addition to RNN, we adopted an architecture known as “Transformer”, which can process input signals all at once through parallel processing, which ultimately leads to a more efficient learning of long-term dependencies (Subakan et al., 2021). Transformer has gained competitive performance and considerable popularity in speech recognition (Karita et al., 2019), speech synthesis (Li et al., 2019), speech enhancement (Kim et al., 2020), and audio source separation (Subakan et al., 2021), as well as other applications such as ChatGpt. We adopted the SepFormer from Subakan et al. (2021), a Transformer-based, top-performing model in speech separation applications at the time of this study, according to Papers with Code website. This state-of-the-art SepFormer model was used here as a reference for the flagship benchmark model to explore the maximum benefit/upper bound of DNN-based noise reduction for CIs, while the RNN model served as a relatively low-complexity, but still an advanced model. In addition to including more sophisticated models to assess the potential of DNN solutions and training the models with a large custom-created dataset, the effectiveness of the models was evaluated not only with objective intelligibility metrics, but also with CI listeners. Most studies have given priority to intelligibility. However, it is also important to evaluate the perceived quality. In this work, we evaluated both intelligibility and quality to also investigate the impact of the processing algorithms on the subjective quality of the processed speech.

## 2 Results

### 2.1 Objective evaluations

The models produced significant improvements across all objective evaluation metrics. The objective evaluation scores for both RNN and SepFormer models with 340 test samples are shown in Figures 1a-1c. As shown in Figure 1a, RNN processing significantly improved scale-invariant source to distortion ratio (SI-SDR) scores over the unprocessed condition across all SNRs, for both masker types (2-talker babble (TTB) and CCITT (stationary speech-shaped noise) maskers, paired t tests, *p <* 0.0001). The improvements in speech quality and intelligibility introduced by the RNN model can also be predicted by the other two metrics: “perceptual” evaluation of speech quality (PESQ) (Figure 1b) and short-time objective intelligibility (STOI) (Figure 1c). These improvements are also statistically significant (paired t tests, *p <* 0.0001). Although statistically significant, the improvement in the speech intelligibility metric (i.e., STOI) was not as large as in the two speech quality metrics (i.e., SI-SDR and PESQ). This is probably due to ceiling effects: the SNR tested was high overall (starting from 1 dB SNR) and the predicted speech intelligibility was not a significant issue in these test conditions (Tang et al., 2017). For the SepFormer model, the objective evaluation scores for the processed audio signals had even more separation from the unprocessed baseline, indicating better performance by SepFormer than RNN (paired t tests, *p <* 0.0001 for all three metrics, across all SNRs, and for both masker types). The superior performance of SepFormer over RNN is especially evident in the noisier, 1-dB SNR condition across all three evaluation metrics.

### 2.2 Behavioral testing with CI listeners

#### 2.2.1 Speech-in-noise intelligibility

The speech intelligibility scores were measured behaviorally as percent correct for the following three processing conditions: “unprocessed”, “processed by RNN”, and “processed by SepFormer”. As with the objective evaluation metrics, CI listeners were tested with both TTB and CCITT maskers. The speech intelligibility performance was measured at both 5- and 10-dB SNR test conditions. The DNN models introduced improvements in speech intelligibility scores over the unprocessed condition at both SNRs and with both masker types for almost all CI listeners tested (only a few exceptions occurred mostly with RNN processing, when the masker type was CCITT). To directly reflect the restoration of speech intelligibility, the percent correct scores for all processing conditions (including “unprocessed”) were referenced to the listening scores obtained in quiet without interference (therefore mostly negative values below the zero line, indicating the expected deleterious effect of noise on speech comprehension; referenced as delta percent scores). A series of models with different combinations of the fixed effects (i.e., SNR, processing condition, and masker type) and with subject intercept being a random effect, were manually fitted and compared based on Akaike information criterion (AIC). The best-fit is achieved by modeling the delta percent scores with SNR, processing condition, masker type, and the interaction between processing condition and masker type as fixed effects, with random effects to account for the variability associated with subject IDs: *model* = *lmer*(*delta percent correct ∼ SNR* + *processing condition* + *masker type* + *processing condition* : *masker type* + 1*|subject*). There was a significant main effect of SNR (*F* [1] = 125.91*, p <* 0.0001), processing condition (*F* [2] = 109.52*, p <* 0.0001), masker type (*F* [1] = 9.24*, p* = 0.0028), as well as interaction between processing condition and masker type (*F* [1] = 3.78*, p* = 0.025). The homoscedasticity assumption of the model was checked and validated by plotting the model residuals against the fitted values, and the sample Quantiles against theoretical Quantiles.

**Figure 1.**
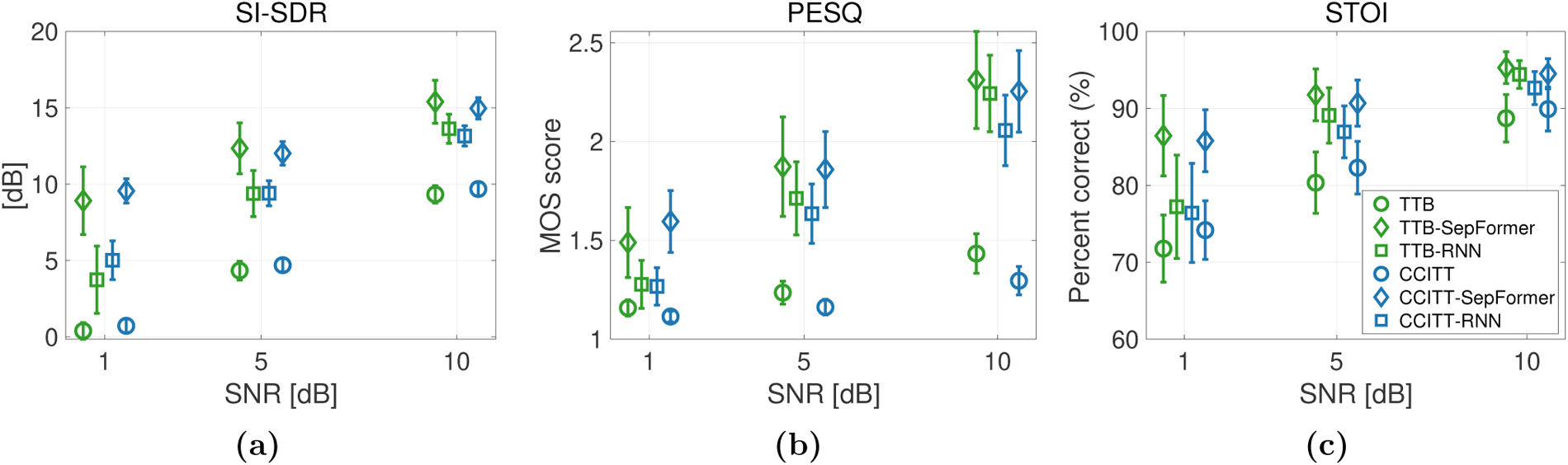
(a)-(c): objective evaluation scores for unprocessed baseline and signals processed with DNN models. Higher scores are better. Statistical significance was found for all cases between the processed and unprocessed conditions, as well as between two processed conditions. MOS: mean opinion scores.

Both the RNN and SepFormer models restored speech intelligibility to performance levels approaching those of quiet settings (i.e., zero line) for both non-speech (Figure 2, left—CCITT) and speech (Figure 2, right—TTB) masker types, across both SNR conditions. Estimated marginal means (EMMs) were computed for all factor level combinations of SNR, processing condition, and masker type using the “emmeans” package in R. The EMMs with standard error and confidence interval for the three processing conditions within each Masker Type and SNR condition are shown in Table 1.

With CCITT masker, post-hoc pairwise comparisons revealed that the scores from RNN (EMM difference (diff) = 11.94, *p < .*0001) and SepFormer (diff = 21.27, *p < .*0001) processing condition were significantly higher than those from unprocessed conditions under the 5-dB SNR configuration. SepFormer also outperformed RNN by 9.32 on average (*p* = .0005). Although the baseline unprocessed condition improved when the SNR was increased to 10 dB (diff = 15.2, *p < .*0001), both RNN and SepFormer continued to improve speech intelligibility scores, with SepFormer still outperforming RNN (diff = 9.32, *p* = 0.005).

Aforementioned benefits from RNN and SepFormer processing over unprocessed conditions, with SepFormer outperforming RNN, are also evident with TTB masker type. However, TTB masker worsened speech intelligibility significantly for the baseline unprocessed condition (diff = 9.03, *p* = 0.0002), reflecting more deleterious impact from speech noise on speech intelligibility than non-speech noise. But both RNN and SepFormer models still improved the speech intelligibility to similar levels as in CCITT-masker conditions. There is no significant difference in speech intelligibility score between TTB and CCITT conditions (RNN: diff = 0.0231, *p* = 0.99; SepFormer: diff = -3.381, *p* = 0.16). In other words, both RNN and SepFormer models introduced *greater* speech enhancement on average when the masker was *speech* over when the masker being non-speech stationary noise.

##### Comparison with other studies

Figure 3 shows the comparison of percent correct improvements introduced by the models in current study and those from several previous studies. In non-speech stationary noise (Figure 3a), models from current studies achieved comparable performance to traditional, signal processing algorithms (Cohen, 2003; Yang & Fu, 2005; Mauger, Arora, & Dawson, 2012). The benefit from those traditional signal processing algorithms is limited in speech babble noise (Figure 3b, Yang & Fu (2005) – 6-talker babble; Mauger, Arora, & Dawson (2012) – 4-talker babble). However, both RNN and SepFormer models introduced even *<u>more</u>* improvements in a 2-talker babble noise compared to steady-state noise, indicating a substantial advantage of deep-learning based noise reduction algorithms in reducing non-stationary, multi-talker noise interference.

**Table 1.** EMMs for processing conditions with different masker type and SNR configurations. SE: standard error; CI: confidence interval.

| Masker type | SNR | Processing condition | EMMs | SE | 95% CI |
| --- | --- | --- | --- | --- | --- |
| CCITT | 5 | Unprocessed | -37.88 | 3.11 | [-44.24, -31.51] |
| CCITT | 5 | RNN | -25.93 | 3.11 | [-32.30, -19.57] |
| CCITT | 5 | SepFormer | -16.61 | 3.11 | [-22.97, -10.25] |
| CCITT | 10 | Unprocessed | -22.63 | 3.11 | [-28.99, -16.27] |
| CCITT | 10 | RNN | -10.69 | 3.11 | [-17.05, -4.33] |
| CCITT | 10 | SepFormer | -1.37 | 3.11 | [-7.73, 5.00] |
| TTB | 5 | Unprocessed | -46.91 | 3.11 | [-53.27, -40.54] |
| TTB | 5 | RNN | -25.91 | 3.11 | [-32.27, -19.55] |
| TTB | 5 | SepFormer | -19.99 | 3.11 | [-26.35, -13.63] |
| TTB | 10 | Unprocessed | -31.66 | 3.11 | [-38.02, -25.30] |
| TTB | 10 | RNN | -10.67 | 3.11 | [-17.03, -4.30] |
| TTB | 10 | SepFormer | -4.75 | 3.11 | [-11.11, 1.62] |

#### 2.2.2 Subjective evaluation of the quality

In addition, as shown in Figure 4, participants rated the quality of the processed speech by both RNN and SepFormer models in both TTB and CCITT maskers as similar to that of the unprocessed signals before mixing with noise (i.e., in quiet). A two-way analysis of variance (ANOVA, with repeated measures) was carried out, involving the processing condition (quite, RNN, SepFormer), the type of masker (TTB, CCITT) as within-subject factors. The test results indicated that there were no statistically significant effect of the processing condition (*F* [2] = 1.64*, p* = 0.2), or the type of masker (*F* [1] = 0.58*, p* = 0.45), or two-way interaction between the processing condition and the type of masker (*F* [2] = 0.57*, p* = 0.57). This suggests that model processing did not significantly distort the speech quality while suppressing the background interference.

#### 2.2.3 Improvement with models vs. demographic factors

We performed a linear mixed effects model with the change in score relative to the unprocessed noisy condition as the dependent variable, masker type (TTB/CCITT), model (RNN/SepFormer), and demographic factors as independent variables, and ID as a random factor. The demographic information is listed in Table 2, including gender, age at testing, duration of CI use, age at onset of hearing loss, number of active electrodes, and coding strategy. A stepwise regression analysis was performed on the full linear model, and the output of the stepwise analysis was a linear model that only included masker type and DNN model, with ID as a random factor. Demographic factors always increased the Akaiki criterion of the linear model. This suggests that the potential benefit of the DNN-based speech enhancement algorithms that has been shown so far was not related to any of the selected demographic factors. The data sample might not be powered sufficiently to establish relationships between benefits from models and demographic factors. However, considering the large variability that is typical of CI population, these results, for this sample, demonstrate the promise of DNN models towards clinical application in the near future for better noise reduction in more complex listening environments (e.g., with multi-talker noise interference).

**Figure 2.**
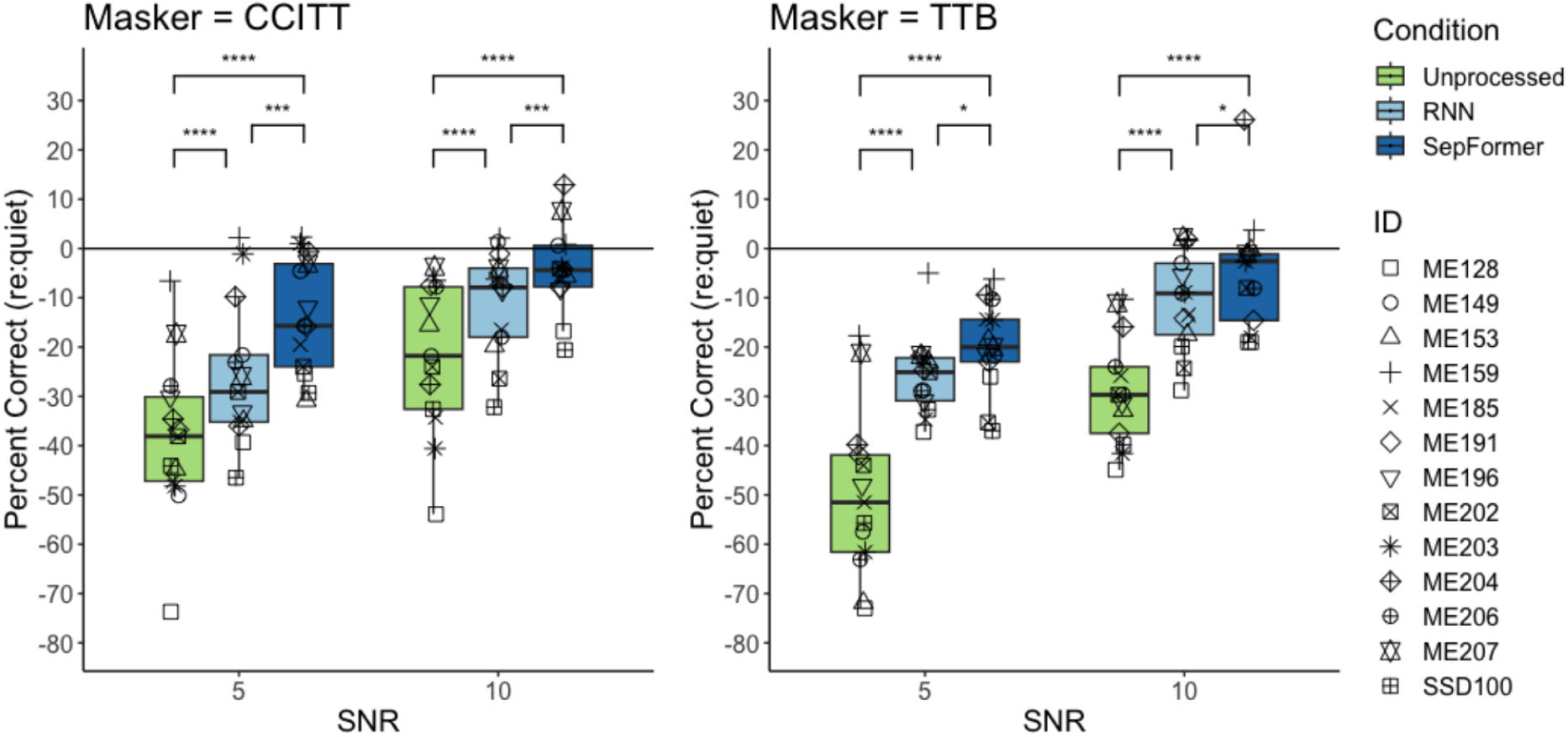
Individual sentence recognition performance (referenced to performance in quiet) plotted as a function of each processing condition for both SNR conditions, separately CCITT and TTB maskers. The boxes depict the values between the 25th and 75th percentiles, and the whiskers represent minimum and maximum values. Medians are shown as horizontal lines. Significance stars: .05 > ∗ ≥ .01, .001 > ∗ ∗ ∗ ≥ .0001, .0001 > ∗ ∗ ∗∗.

## 3 Discussion

In this work, we implemented, trained, and tested the RNN and SepFormer models for noise reduction in CIs. The models were evaluated with objective metrics and behaviorally with a total of 13 CI listeners. Compared to widely adopted classification and regression models such as convolutional neural network (CNN) models, RNN does not require an input of fixed dimensions such as an image file with standardized size, which makes it more suitable for processing audio samples of varying lengths. RNN also keeps a memory of prior information and makes predictions based on both previous and current information (i.e., through LSTM). This feature is important for input signals such as speech, of which segments of information across time interconnect with each other to form the contextual meaning. SepFormer is a transformer-based model that takes the step further by extending RNN’s sequential processing to parallel processing, which best suits the need for processing longer sequences of signals. These two models were then trained with a large custom dataset containing various target-noise configurations, totaling 22360 instances for each training iteration. More specifically, we created the large training and validation dataset from commonly used speech and non-speech corpora in machine-listening field—LibriSpeech and WHAM!, respectively. The IEEE corpus, which is more common in hearing research, and CCITT noise were used for creating the testing dataset. The models were also trained more extensively with 100 learning cycles, to explore the full capacity of the models in noise reduction performance.

**Figure 3.**
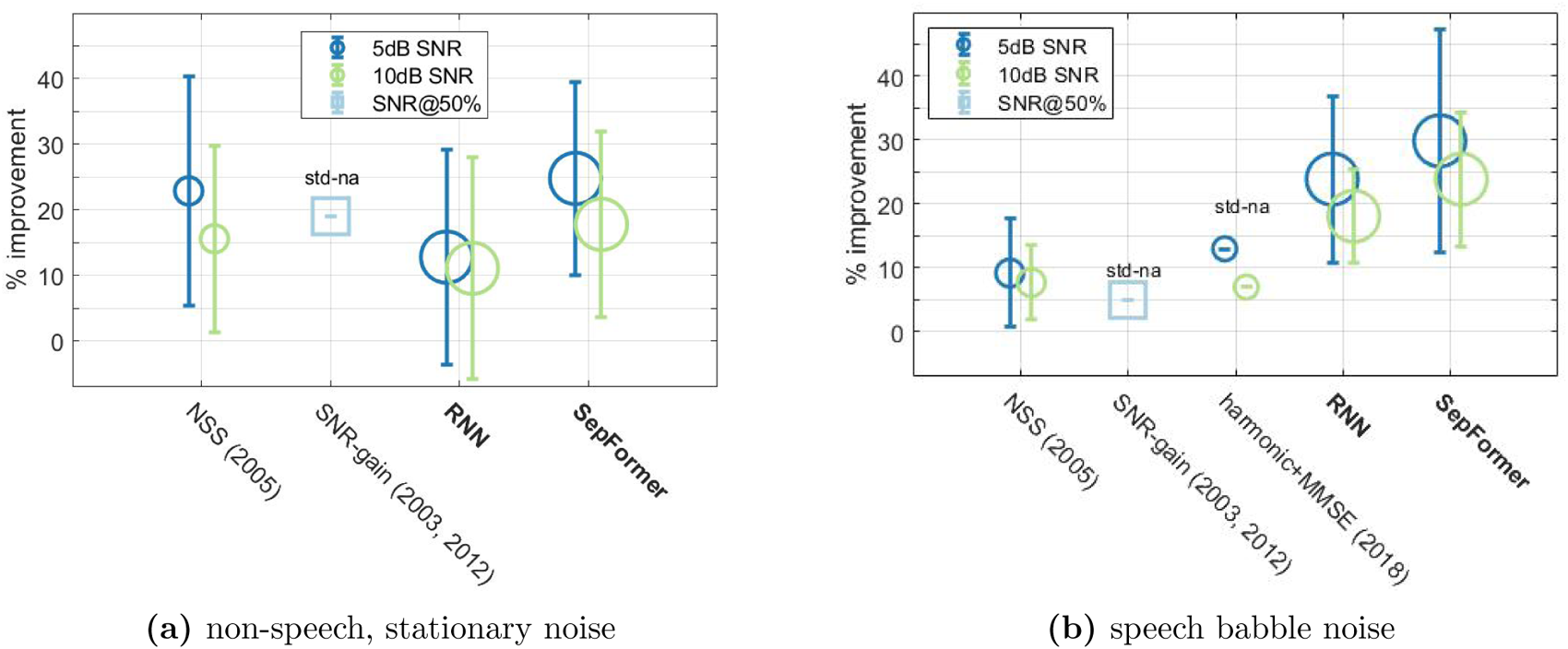
Comparison of model performance between current study and previous studies tested in similar conditions. The size of the circle is proportional to the number of CI participants tested in the study. The center of the circles align with population mean and the error bar is 1 standard deviation. Note that values from some studies are visual approximation from the figures, which explains the lack of error bar (indicated by text annotation: “std-na”). Except for “harmonic+MMSE (2018)”, all models listed from other studies are traditional signal processing algorithms that are not machine-learning based.

**Figure 4.**
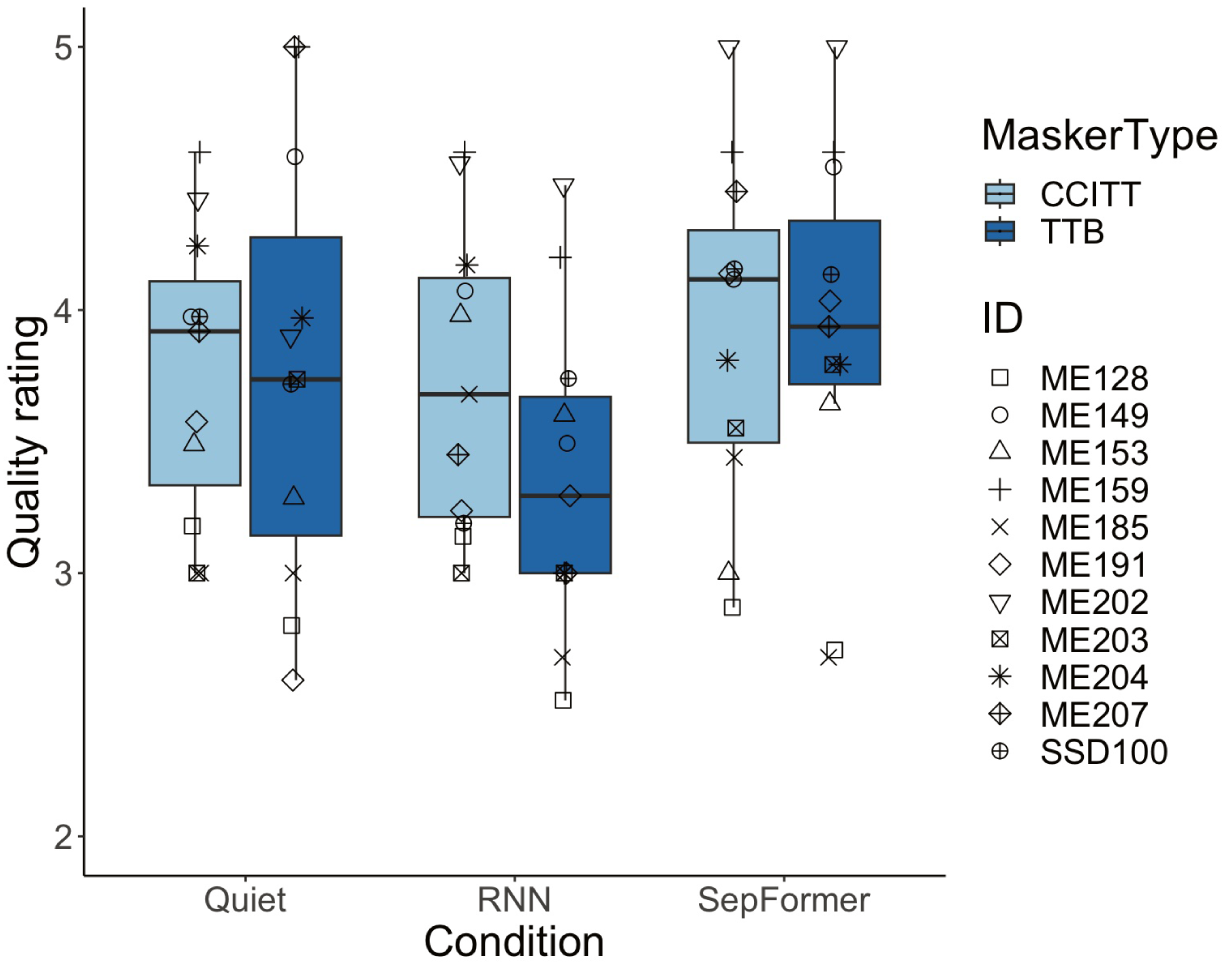
Subjective evaluation scores, according to the mean opinion score (MOS) scale. The score assigns a numerical measure of the human-judged overall quality: 1 corresponds to “bad”, while 5 represents “excellent”. Two blocks of testing were conducted for the “Quiet” condition to equalize the number of samples since there were two types of maskers for each processing condition. Note that the quality was evaluated for two processing conditions in 10-dB SNR. Due to time restriction, the quality was not evaluated for 5-dB or unprocessed conditions.

Both RNN and SepFormer models significantly improved CI listener’s speech intelligibility in noise relative to unprocessed, noisy speech (almost to the levels in quiet in certain conditions). The improvement in speech intelligibility in stationary non-speech noise was comparable to those gains achieved by traditional signal processing strategies. More importantly, both RNN and SepFormer introduced substantially ***more*** improvements in non-stationary ***multi-talker*** noise interference, where conventional signal processing strategies are limited, than stationary noise.

The objective evaluation metrics did not predict the observed advantage of the two DNN models in multi-talker background noise, which underscores the importance of testing the models with CI participants. It is also noteworthy that the improvements in intelligibility did not come at the price of compromised sound quality of the speech among CI listeners. The fact that there were no significant contributions from any demographic disparity across participants suggests that the algorithms have the potential to broadly benefit the general CI population regardless of their hearing etiology, duration of deafness, experience with CI listening, and processor settings; although, it should be noted that the predictive power of the model was limited by the number of participants. Nevertheless, these results show the promise of using machine-learning based models as a complementary or even replacement algorithm for current existing signal processing strategies, to tackle more complex listening challenges involving multi-talker noise interference.

Given that SepFormer was the top-performing, state-of-the-art model for speech separation and enhancement at the time of the study, while RNN is a basic two-layer template model, it is not surprising that the SepFormer outperformed RNN in every scenario. It provided an estimate of the current upper limit to which a DNN-based strategy could restore speech comprehension when paired with CI devices. However, SepFormer is a very complicated model, containing over 26 million parameters. The processing time of such a computationally heavy model turned out to be almost 5 times the duration of the incoming signal on average, which renders it not suitable for a real-time applications such as a CI audio processor. Note that the inference and calculation of the processing time was conducted on CPU (MacBook Pro i7 2.2GHz) instead of GPU, considering hearing aids and CIs mostly adopt DSPs (digital signal processors). The time constraint for the processing delay in a real-time device such as CI should be below about 10-20 ms to avoid disturbance in speech production and audio-visual integration (Stone & Moore, 1999; Goehring et al., 2018, 2019; Bramsløw et al., 2018). This timing delay should be even lower for individuals with single sided deafness who are fitted with CI on the side with hearing deprivation while receiving acoustic input on the other side (Zirn et al., 2015). Another limitation comes from the high demand for computational power and memory. Thus, at this time, it is probably unrealistic to execute these highly complex state-of-the-art models even in high-end audio DSPs used in the CI processors. The RNN model, on the other hand, is a much computationally lighter model and the processing only took 3% of the input signal duration on average. This implies that RNNs can in theory retain low-latency inference, which makes them better-suited DNN model for real-time speech enhancement in CIs. At the time of this study, Oticon launched the world’s first hearing aid with an on-board DNN—Oticon More, which is an encouraging message for the field working on the implementation of DNN-based noise-reduction solutions for CIs.

Future plans include optimizing and lightening the DNN models and testing them real-time in research processors with CI listeners. The training and testing should also be expended to languages other than English. Finally, the deployment of the DNN models in research processors should be tested against existing commercial solutions for noise reduction, as well as be further evaluated in a wide variety of realistic listening environments outside research labs.

## 4 Methods

### 4.1 Participants

A total of thirteen adults (seven males) fitted with MED-EL CIs (MED-EL GmbH, Innsbruck, Austria) participated in the study. They were between the ages of 20 and 72 (mean = 58.6 years, SD = 14.7 years). The average duration of CI use was 6.5 years (SD = 5 years). Demographic information is detailed in Table 2, including each subject’s default clinical sound coding strategy, which was used when listening to test materials. This study was approved by the Western Institutional Review Board (Protocol 20100066). All subjects gave informed written consent prior to testing. Only research subjects whose participation in the study would cause financial hardship received financial compensation for their participation.

### 4.2 Deep Neural Network Architectures

#### 4.2.1 RNN

The schematic diagram of the single-channel, RNN-based speech enhancement algorithm is illustrated in Figure 5a. A clean target speech signal and either speech babble or non-speech noise were mixed to create unprocessed noisy speech. The features used as input to the RNN model were the spectral magnitudes of the short-time Fourier transformation (STFT) of the mixtures (size of Fast Fourier Transform = 512). The spectral magnitudes were extracted using Hamming-windowed frames with a window size of 32 samples and a hop size of 16 samples applied to signals sampled at 16 kHz. “Add-one” log (i.e., adding 1 to the value before taking log) was applied to the spectral magnitude to reduce the influence of values that are smaller than 1. The predicted mask (i.e., the model outcome) was applied to the STFT of the mixture (spectral magnitudes) to generate the “de-noised” spectrum of the mixture and was a continuous “soft” mask (continous gains from 0 to 1, as opposed to a binary mask where the values of the mask are either one or zero). Previous work suggests that “soft” continuous masks result in better speech quality and intelligibility than binary masks under various noise conditions (Madhu et al., 2013). The “de-noised” estimate would ideally consist of the enhanced target speech only. This “de-noised” spectrum was compared with the spectrum of the clean target speech to compute the mean square error (MSE) loss for training optimization. The processed speech estimate was then recovered by resynthesizing (i.e., taking the inverse STFT) the “de-noised” spectrum.

The RNN network consisted of an input layer (size = 512) and two hidden LSTM layers (256 units), with each LSTM followed by a projection layer (128 units). A rectified linear unit (ReLU) was used as an activation function. A PyTorch-powered speech toolkit—SpeechBrain—was used to implement, train, and test the RNN model. The Adam optimizer was used for minimizing the MSE loss during the training process (Kingma & Ba, 2017), with learning rate set at 0.0001. The model performance was evaluated and monitored with a validation dataset at the end of each learning cycle (i.e., one epoch, containing all training samples). The training was terminated after 100 epochs as the model performance with validation dataset stabilized with no further substantial improvements.

**Table 2.** Demogcraphic information for the subjects who participated in this study. Age_T: age at the time of testing (years); Age_HL: age of onset of hearing loss (years); Dur CI: duration of CI use (months); #_El: number of active electrodes; Score: IEEE score in quiet (percent correct). Strategy: 1 – Fine structure processing with *parallel* stimulation in the four apical channels; 2 – Fine structure processing with *sequential* stimulation in the four apical channels; 3-Fine structure processing with *sequential* stimulation in a variable number of apical channels

| Subject | Gender | Age_T | Age_HL | Dur_CI | #_El | Strategy | Score |
| --- | --- | --- | --- | --- | --- | --- | --- |
| ME128 | M | 70 | 30 | 95 | 10 | FS4-p<sup>1</sup> | 79 |
| ME149 | F | 57 | 27 | 53 | 8 | FS4<sup>2</sup> | 92 |
| ME153 | M | 71 | 34 | 45 | 12 | FS4 | 91 |
| ME159 | F | 60 | 17 | 64 | 12 | FSP<sup>3</sup> | 92 |
| ME185 | F | 65 | 8 | 119 | 12 | FS4-p | 76 |
| ME191 | M | 64 | 42 | 40 | 11 | FS4-p | 93 |
| ME196 | M | 65 | 11 | 24 | 11 | FS4 | 92 |
| ME202 | M | 55 | 0 | 40 | 12 | FS4 | 56 |
| ME203 | F | 63 | 5 | 41 | 12 | FS4-p | 73 |
| ME204 | M | 63 | 41 | 33 | 12 | FS4-p | 48 |
| ME206 | F | 72 | 27 | 66 | 12 | FS4-p | 92 |
| ME207 | M | 20 | 0 | 229 | 12 | FS4 | 86 |
| SSD100 | F | 37 | 24 | 158 | 12 | FS4-p | 68 |

#### 4.2.2 SepFormer

While the light-weight, generic RNN model serves as a proof of concept for DNN algorithms that are suitable for real-time processing in hearing devices such as CIs, we also implemented the current state-of-the-art model for speech separation applications—SepFormer, to explore the ceiling limit of current DNN technology in noise reduction for CIs (especially non-stationary, multi-talker noise interference). The model architecture is shown in Figure 5b. Instead of directly feeding STFT features of the noisy mixture to the network as in the RNN model, a single-layer convolutional network was used as an encoder to learn the 2-dimensional features of the input noisy signal (256 convolutional filters with a kernel size of 16 samples and a stride factor of 8 samples). Similarly, at the end of the process, a transposed convolution layer with the same stride and kernel size as in the encoder was used to turn the source-separated features into separate sources. The extracted features of the noisy mixture go into the masking network, which estimates the masks for the foreground (i.e., target speech) and background. These masks were also continuous or soft-decision masks that provided continuous gains from 0 to 1 as in the RNN model. In the masking network, the features were first normalized and processed by a linear layer. They were then buffered into chunks of size 250 along the time axis with an overlap factor of 50%. Next, they were fed into the core of the masking net—SepFormer block. This block consists of two transformer structures that learned both short and long-term dependencies. More details about this model can be found in Subakan et al. (2021). The output of the SepFormer block was then processed by a parametric rectified linear unit (PReLU) and linear layer. The overlap-add scheme, described in Luo & Mesgarani (2019), was used to sum up the chunks. This summed representation was passed through two feed-forward layers and a ReLU activation function to finally generate the masks for both the foreground and background sources. The training procedure and infrastructure were the same as for the RNN model.

**Figure 5.**
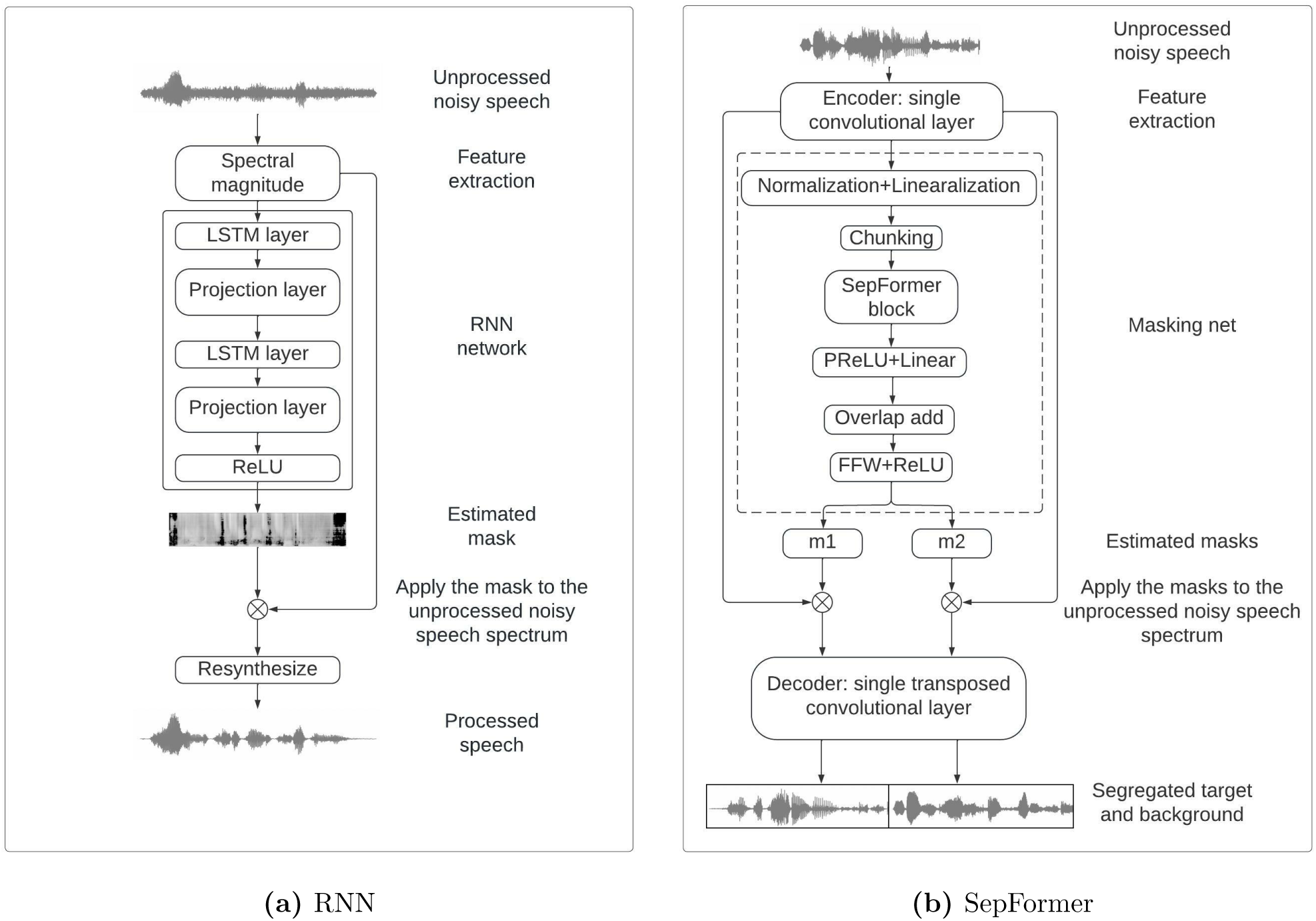
Schematic diagrams of the DNN architecture and signal processing frameworks used in this study: (a) RNN and (b) SepFormer

### 4.3 Dataset

#### 4.3.1 Training and validation datasets

Clean target speech audio signals were from LibriSpeech ASR corpus, an open-source, large-scale (*∼*1000 hours, 2484 speakers) corpus of read English speech. The speech audio samples were extracted from read audiobooks from the LibriVox initiative, and every recording was carefully segmented and aligned with one another. This study used the training subset that contains 100-hours of clean speech. The non-speech materials were from WHAM!, an open-source, large-scale (*∼* 82 hours) dataset of noise audio signals. The noise data samples were recorded from different urban spots across the San Francisco Bay Area, mainly comprising restaurants, cafes, bars, and parks. Each recording was processed to eliminate any parts that include intelligible speech. Four types of noisy speech were created for training: target in non-speech noise (WHAM! noise), and target in 1, 2, 4-talker babble. Recordings from LibriSpeech were mixed to form multi-talker babble. All speakers in the mixture are distinct from one another and speaking different content. Each type of noisy speech was mixed at SNRs from 1 to 10 dB in 1-dB steps, with equal representation. The loudness of target was kept constant at ITU level of 29. A total of 5590 mixtures were created for each noise type, which resulted in a training dataset of *∼* 30 hours. The validation dataset contains 410 mixtures for each type of noisy speech. To speed up the process of model preparation, the training and validation were conducted on graphics processing unit (GPU, Google Colaboratory).

#### 4.3.2 Testing dataset

Clean target speech materials were extracted from IEEE corpus, which is often used in auditory research and clinics. This corpus contains recordings of 33 different talkers. Most speakers read the full set of 720 IEEE sentences. Three hundred and forty sentences from the IEEE corpus were mixed with CCITT noise (speech shaped stationary noise according to ITU-T Rec. G.227) or 2-talker babble at SNRs of +1, +5 and +10 dB. Two-talker babbles were produced by mixing the IEEE recordings from 120 sentences that were not chosen as target sentences. Male recordings were used for target while female recordings were used to generate 2-talker babbles to create the noisy mixture. Note that IEEE corpus was never used during training or validating the models. The non-speech noise used for training and validation was relatively sparse environmental sounds, whereas the CCITT used for testing was spectral-temporally dense stationary noise. This testing dataset was used for both objective evaluations (section 4.4.1) and CI subject testing (section 4.4.2). Two hundred and froty sentences from the remaining IEEE sentences were used for subject testing in quiet. The model testing was conducted on a MacBook Pro, i7 core (2.2GHz) central processing unit (CPU).

### 4.4 Procedure

#### 4.4.1 Objective evaluations

The DNN models were first evaluated quantitatively using three commonly used acoustic evaluation metrics: scale-invariant source-to-distortion ratio (SI-SDR) (Roux et al., 2018), short-time objective intelligibility (STOI) (Taal et al., 2011, 2010), and “perceptual” evaluation of speech quality (PESQ) (Rix et al., 2001; Hu & Loizou, 2008a). These objective evaluation methods helped inform the overall expected benefit before conducting behavioral listening test with CI users. All three evaluation metrics compare the clean reference speech and the same speech recovered from the noisy mixture and quantify the agreement between the two, which allows for an estimate of the improvement in quality and intelligibility due to model processing when compared to the metric for the unprocessed noisy mixture.

The traditional SDR metric decomposes the estimated source into four components representing respectively the true source, spatial distortions, interference, and artifacts. The final SDR score is computed by calculating the ratio of the source energy to the sum of all other projection energies (i.e., spatial distortions, interference, and artifacts) as described in Vincent et al. (2006). The SI-SDR with slight modifications as described in Roux et al. (2018) has been shown to be more robust and is now the standard for DNN noise reduction evaluation.

The STOI metric was initially designed to predict the intelligibility of speech processed by enhancement algorithms. Recently, Falk et al. (2015) demonstrated that STOI outperformed all other measures for predicting intelligibility of CI listeners. The STOI first applies time-frequency analysis to both clean reference and processed speech signals. An intermediate intelligibility measure is obtained by estimating the linear correlation coefficient between clean and processed time-frequency units. The final STOI score is the average of all intermediate intelligibility estimates from all time-frequency units.

The PESQ score ranges between –0.5 and 4.5. It was calculated by comparing the reference signal with the processed signal by deploying a perceptual model of the human auditory system. The PESQ is computed as a linear combination of average disturbance value and average asymmetric disturbance value. The parameters for the linear combination can be further modified towards predicting different aspects of speech quality. More details can be found in Rix et al. (2001), Hu & Loizou (2008a) and Kokkinakis & Loizou (2011). In general, the PESQ has been shown to be capable of reliably predicting the quality of processed speech. Kokkinakis & Stohl (2021) showed that parameter optimization for reverberation suppression algorithms based on the PESQ metric resulted in better performance than the STOI metric. Therefore, in the present context, the PESQ was chosen to detect and quantify the overall effects of DNN processing on the signal quality.

Note that these objective evaluation metrics were developed based on data from NH listeners, and in this study, these metrics were calculated to mainly inform rather than replace the subject testing. Even the models adapted for CI listening can never fully approximate the actual testing with CI listeners due to the many sources of variability in CI outcomes (Blamey et al., 2009, 2012).

#### 4.4.2 Behavioral testing

##### Test setup

All participants were tested using their everyday clinical program, as listed in Table 2. The stimulus was delivered to each subject’s own audio processor through a direct audio input (DAI) cable, which attenuates the microphone inputs by approximately 30 dB relative to the direct input signal. The DAI also bypasses the front-end directionality and wind-noise reduction features. The test stimuli were presented at an input level corresponding to 65 dB SPL (root mean square (RMS) level). None of the participating subjects used MED-EL’s channel-specific, ambient and transient noise reduction algorithms in their daily maps at the time of testing. At the beginning of the testing session, the audiologist provided instructions regarding the study procedures, and then connected the recipient’s processor to the audio port of a Windows-based touchscreen tablet (Microsoft Surface Pro) through the DAI cable. The proprietary psychophysical software suite, PsyWorks v.6.1 (MED-EL GmbH, Innsbruck, Austria), was used to present the speech stimuli from the tablet to the audio processor. The calibration was performed using a built-in feature within the PsyWorks software and was adjusted according to each recipient’s audio processor type.

##### Intelligibility and quality measurements

Each processing condition (unprocessed, RNN, SepFormer) was evaluated with a list of twenty sentences from the test dataset for each combination of masker type and SNR (+10 and +5 dB SNR). Each subject performed a total of 13 tests (2 masker types x 3 processing conditions x 2 SNRs + 1 quiet). The testing was carried out in a self-administered manner. The subjects used the tablet and PsyWorks to present the speech materials to their own audio processors. Subjects were assigned a unique presentation order using a Latin square design and were blinded to the processing condition. The subjects either vocalized their responses through a microphone located in front of them or typed them, according to their preference. The responses were captured in real-time by an automatic speech-to-text module (Google API) in the case of spoken responses. Spoken responses could be edited by typing before submission, and the PsyWorks software automatically scored words as correctly or incorrectly identified. Words containing additions, substitutions, or omissions were scored as incorrect. The percent correct scores for each condition were calculated by dividing the number of correct words by the total number of words. After each list, the percent correct was displayed and stored electronically. All participants were native English speakers, and none of the participants that elected to speak their responses had speech difficulties that jeopardized the scoring of their responses. The total testing time for all experimental conditions tested was approximately 2.5 hours including multiple breaks.

## 5 Author Declarations

### Funding Acknowledgements

This research was supported by funding from the National Institutes of Health [Grant R01DC015989]; and MED-EL Corporation.

### Ethics Approval statement

This study was approved by the Western Institutional Review Board (Protocol 20100066). All subjects gave informed written consent prior to testing. Research subjects whose participation in the study would have caused financial hardship received financial compensation for their participation.

### Data Accessibility

Data will be made available upon publication.

## Acknowledgements

We thank our research subjects for their participation in this study. We also thank Jenna Felder for helping collect data from CI participants.

